# Orienting the causal relationship between imprecisely measured traits using genetic instruments

**DOI:** 10.1101/117101

**Authors:** Gibran Hemani, Kate Tilling, George Davey Smith

## Abstract

Inference of the causal structure that induces correlations between two traits can be achieved by combining genetic associations with a mediation-based approach, as is done in the causal inference test (CIT) and others. However, we show that measurement error in the phenotypes can lead to mediation-based approaches inferring the wrong causal direction, and that increasing sample sizes has the adverse effect of increasing confidence in the wrong answer. Here we introduce an extension to Mendelian randomisation, a method that uses genetic associations in an instrumentation framework, that enables inference of the causal direction between traits, with some advantages. First, it is less susceptible to bias in the presence of measurement error; second, it is more statistically efficient; third, it can be performed using only summary level data from genome-wide association studies; and fourth, its sensitivity to measurement error can be evaluated. We apply the method to infer the causal direction between DNA methylation and gene expression levels. Our results demonstrate that, in general, DNA methylation is more likely to be the causal factor, but this result is highly susceptible to bias induced by systematic differences in measurement error between the platforms. We emphasise that, where possible, implementing MR and appropriate sensitivity analyses alongside other approaches such as CIT is important to triangulate reliable conclusions about causality.

## Introduction

Observational measures of the human phenome are growing ever more abundant, but using these data to make causal inference is notoriously susceptible to many pitfalls, with basic regression-based techniques unable to distinguish a true causal association from reverse causation or confounding (1-3). In response to this, the use of genetic associations to instrument traits has emerged as a technique for improving the reliability of causal inference in observational data, and with the coincident rise in genome-wide association studies it is now a prominent tool that is applied in several different guises (3-6). However, potential pitfalls remain and one that is often neglected is the influence of non-differential measurement error on the reliability of causal inference.

Measurement error is the difference between the measured value of a quantity and its true value. This study focuses specifically on non-differential measurement error where all strata of a measured variable have the same error rate, which can manifest as changes in scale or measurement imprecision (noise). Such variability can arise through a whole plethora of mechanisms, which are often specific to the study design and difficult to avoid (7,8). Array technology is now commonly used to obtain high throughput phenotyping at low cost, but comes with the problem of having imperfect resolution, for instance methylation levels as measured by the Illumina450k chip are prone to have some amount of noise around the true value due to imperfect sensitivity (9,10). Relatedly, if the measurement of biological interest is the methylation level in a T cell, then measurement error of this value can be introduced by using methylation levels from whole blood samples because the measured value will be an assay of many cell types (11).

Measurement error will of course arise in other types of data too. For example when measuring BMI one is typically interested in using this as a proxy for adiposity, but it is clear that the correlation between BMI and underlying adiposity is not perfect (12). A similar problem of biological misspecification is unavoidable in disease diagnosis, and measuring behaviour such as smoking or diet is notoriously difficult to do accurately. Measurement error can also be introduced after the data have been collected, for example the transformation of non-normal data for the purpose of statistical analysis will lead to a new variable that will typically incur both changes in scale and imprecision (noise) compared to the original variable. The sources of measurement error are not limited to this list (8), and its impact has been explored in the epidemiological literature extensively (13,14). Given the near-ubiquitous presence of measurement error in phenomic data it is vital to understand its impact on the tools we use for causal inference.

An established study design that can provide information about causality is randomisation. Given the hypothesis that trait A (henceforth referred to as the exposure) is causally related to trait B (henceforth referred to as the outcome), randomisation can be employed to assess the causal nature of the association by randomly splitting the sample into two groups, subjecting one group to the exposure and treating the other as a control. The association between the exposure and the outcome in this setting provides a robust estimate of the causal relationship. This provides the theoretical basis behind randomised control trials, but in practice randomisation is often difficult or impossible to implement in an experimental context due to cost, scale or inability to manipulate the exposure. The principle, however, can be employed in extant observational data through the use of genetic variants associated with the exposure (instruments), where the inheritance of an allele serves as a random lifetime allocation of differential exposure levels (15,16). Two statistical approaches to exploiting the properties of genetic instruments are widely used: mediation-based approaches and Mendelian randomisation (MR).

Mediation-based approaches employ genetic instruments (typically single nucleotide polymorphisms, SNPs) to orient the causal direction between the exposure and the outcome. If a SNP is associated with an exposure, and the exposure is associated with some outcome, then it logically follows that in this simple three-variable scenario the estimated direct influence of the SNP on the outcome will be zero when conditioning on the exposure. Here, the exposure completely mediates the association between the SNP and the outcome, providing information about the causal influence of the exposure on the outcome. This forms the basis of a number of methods such as genetical genomics (17), the regression-based causal inference test (CIT) (4,18), a structural equation modelling (SEM) implementation in the NEO software (5), and various other methods including Bayesian approaches (6). They have been employed by a number of recent publications that make causal inferences in large scale ‘omics datasets (6,19-23).

MR can be applied to the same data - phenotypic measures of the exposure and the outcome variables and a genetic instrument for the exposure - but the genetic instrument is employed in a subtly different manner. Here the SNP is used as a surrogate for the exposure. Assuming the SNP associates with the outcome only through the exposure, the causal effect of the exposure on the outcome can be estimated by scaling the association between the SNP and the outcome by the association between the SNP and the exposure. Though difficult to test empirically, this assumption can be relaxed in various ways when multiple instruments are available for a putative exposure (24,25) and a number of sensitivity tests are now available to improve reliability (26).

By utilising genetic instruments in different ways, mediation-based analysis and MR models have properties that confer some advantages and some disadvantages for reliable causal inference. In the CIT framework (described fully in the Methods) for example, the test statistic is different if you test for the exposure causing the outcome or the outcome causing the exposure, allowing the researcher to infer the direction of causality between two variables by performing the test in both directions and choosing the model with the strongest evidence. The CIT also has the valuable property of being able to distinguish between several putative causal graphs that link the traits with the SNP (Figure 1). Such is not the case for MR, where in order to infer the direction of causality between two traits the instrument must have its most proximal link with the exposure, associating with the outcome only through the exposure.

**Figure 1.**
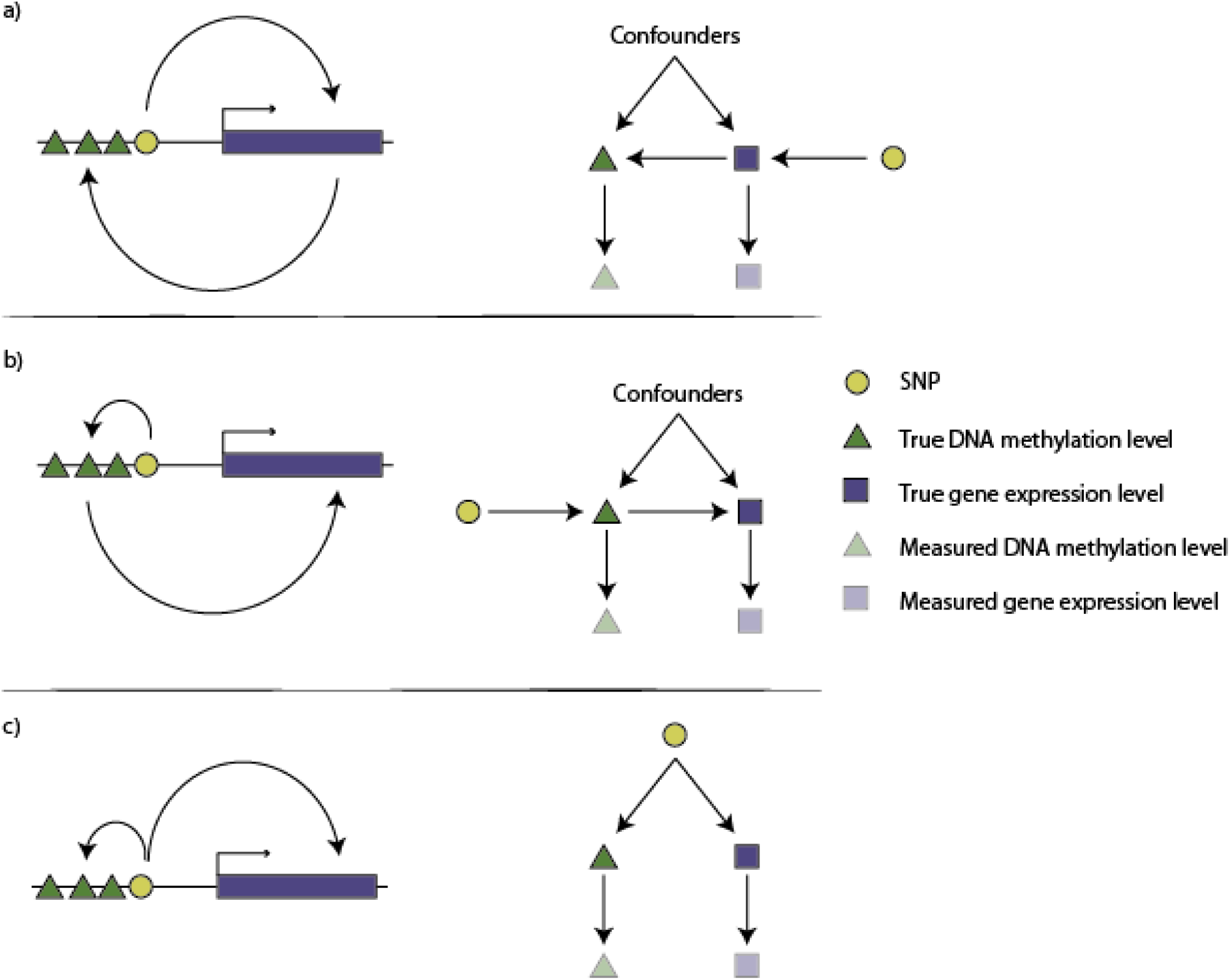
Gene expression levels (blue blocks) and DNA methylation levels (green triangles) may be correlated but the causal structure is unknown. If a SNP (yellow circle) is associated with both DNA methylation and gene expression levels then it can be used as an instrument, but there are three basic competing models for these variables. The causal inference test (CIT) attempts to distinguish between them. a) Methylation causes gene expression. The left figure shows that the SNP influences methylation levels that in turn influence gene expression levels. The right figure shows the directed acyclic graph that represents this model. Faded symbols represent the measured values whereas solid symbols represent the true values. b) The same as in A, except the causal direction is from gene expression to DNA methylation. c) A model of confounding, where gene expression and DNA methylation are not causally related, but the SNP influences them each through separate pathways or a confounder.

Assuming biological knowledge of genetic associations can be problematic because if there exists a putative association between two variables, with the SNP being robustly associated with each, it can be difficult to determine which of the two variables is subject to the primary effect of the SNP (i.e. for which of the two variables is the SNP a valid instrument? Figure 1). By definition, we expect that if the association is causal then a SNP for the exposure will be associated with the outcome, such that if the researcher erroneously uses the SNP as an instrument for the outcome then they are likely to see an apparently robust causal association of outcome on exposure. Genome-wide association studies (GWASs) that identify genetic associations for complex traits are, by design, hypothesis free and agnostic of genomic function, and it often takes years of follow up studies to understand the biological nature of a putative GWAS hit (27). If multiple instruments are available for an hypothesised exposure, which is increasingly typical for complex traits that are analysed in large GWAS consortia, then techniques can be applied to mitigate these issues (16). But these techniques cannot always be applied in the case of determining causal directions between 'omic measures where typically only one cis-acting SNP is known. For example if a DNA methylation probe is associated with expression of an adjacent gene, then is a cis-acting SNP an instrument for the DNA methylation level, or the gene expression level (Figure 1)?

MR has some important advantages over the mediation-based approaches. First, the mediation-based approaches require that the exposure, outcome and instrumental variables are all measured in the same data, whereas recent extensions to MR circumvent this requirement, allowing causal inference to be drawn when exposure variables and outcome variables are measured in different samples (28). This has the crucial advantage of improving statistical power by allowing analysis in much larger sample sizes, and dramatically expands the breadth of possible phenotypic relationships that can be evaluated (26). Second, the mediation-based approach of adjusting the outcome for the exposure to nullify the association between the SNP and the outcome is affected by unmeasured confounding of the exposure and outcome. This is because adjusting the outcome by the exposure induces a collider effect between the SNP and outcome (29), and the in order to fully abrogate this association one must also adjust for all (hidden or otherwise) confounders. MR does not suffer from this problem because it does not test for association through adjustment. Third, when MR assumptions are satisfied the method is robust to there being measurement error in the exposure variable (30). Indeed instrumental variable (IV) analysis was in part initially introduced as a correction for measurement error in the exposure (31), whereas it has been noted that both classic mediation-based analyses (13,14,32,33) and mediation-based methods that use instrumental variables (34,35) are prone to be unreliable in its presence.

Using theory and simulations we show how non-differential measurement error in phenotypes can lead to unreliable causal inference in the mediation-based CIT method. We then present an extension to MR that allows researchers to ascertain the causal direction of an association even when the biology of the instruments are not fully understood, and also a metric to evaluate the sensitivity of the result of this extension to measurement error. Together these extensions improve the utility of MR in cases where mediation based methods might have otherwise been used preferentially. Finally, we apply this method to infer the direction of causation between DNA methylation levels and gene expression levels. Our analyses highlight that because these different causal inference techniques have varying strengths and weaknesses, triangulation of evidence from as many sources as possible should be practiced in causal inference (36).

## Model

We model a system whereby some exposure *x* has a causal influence *β_x_* on an outcome *y* such that

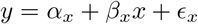

In addition, the exposure is influenced by a SNP *g* with an effect of *β_g_* such that

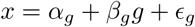

The *α*\* terms represent intercepts, and henceforth can be ignored. The *ϵ_*_* terms denote random error, assumed independently and normally distributed with mean zero. Mediation-based analyses that test whether *x* causally relates to *y* rely on evaluating whether the influence of *g* on *y* can be accounted for by conditioning on *x*, such that

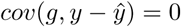

where *ŷ* = *β̂_x_x* and assuming no intercept *y* − *ŷ* = *ϵ_x_*. MR analysis estimates the causal influence of *x* on *y* by using the instrument as a proxy for *x*, such that

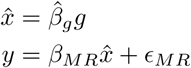

where *β_MR_* ≠ 0 denotes the existence of causality, and *β_MR_* is an estimate of the causal effect.

Measurement error of an exposure can be modeled as a transformation of the true value (*x*) that leads to the observed value, *x*_0_ *= f* (*x*). For example, following Pierce and VanderWeele (30) we can define

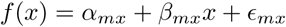

where *α_mx_* and *β_mx_* influence the error in the measurement of *x* by altering its scale, and *ϵ_mx_* represents the imprecision (or noise) in the measurement of *x*. The same model of measurement error can be applied to the outcome variable *y*. In this study we assume there is no measurement error in the SNP, and that measurement error in the exposure and the outcome are uncorrelated.

## Methods

### CIT test

First we describe how the CIT method (4) is implemented in the R package *R/cit* (18). The methodology of the CIT is as follows. Assume an exposure *x* is instrumented by a SNP *g,* and the exposure *x* causes an outcome *y*, as described above. The following tests are then performed:

1. *H*_0_: *cov*(*g,x*) = 0; *H*_1_: *cov*(*g,x*) ≠ 0; *the SNP associates with the exposure*
2. *H*_0_: *cov*(*g, y*) = 0; *H*_1_: *cov*(*g, y*) ≠ 0; *the SNP associates with the outcome*
3. *H*_0_: *cov*(*x, y*) = 0; *H*_1_: *cov*(*x, y*) ≠ 0; *the exposure associates with the outcome*
4. *H*_0_: *cov*(*g, y* − *ŷ*) ≠ 0; *H*_1_: *cov*(*g, y* − *ŷ*) = 0; *the SNP is independent of the outcome when the outcome is adjusted for the exposure*

where *y* − *ŷ* = *y* − *α̂_g_* + *β̂_g_x* is the residual of *y* after adjusting for *x*, where *x* is assumed to mediate the association between the SNP and the outcome. The 4th condition is formulated as an equivalence testing problem that is estimated using simulations, comparing the estimate against from the data against empirically obtained estimates for simulated variables where the independence model is true (full details are given in (4)). We note here that this approach is liable to fail, even when there is a true causal relationship, when confounders of the exposure and outcome are present, as these will induce collider bias.

If all four tests reject the null hypothesis then it is inferred that *x* causes *y*. The CIT measures the strength of causality by generating an omnibus p-value, *p_CIT_*, which is simply the largest (least extreme) p-value of the four tests, the intuition being that causal inference is only as strong as the weakest link in the chain of tests.

Now we describe how we used the CIT method in our simulations. The *cit.cp* function was used to obtain an omnibus p-value. To infer the direction of causality using the CIT method, an omnibus p-value generated by CIT for each of two tests - *p*_*CIT,x*→*y*_, was estimated for the direction of *x* causing *y* (Model 1), and for the direction of *y* causing *x*, *p_CIT_, _y_*_→*x*_ (Model 2). The results from each of these methods can then be used in combination to infer the existance and direction of causality. For some significance threshold *α* there are four possible outcomes from these two tests, and their interpretations are as follows:

- *p_CIT_*,_*x*→*y*_ < *α* and *p*_CIT, *y*→*x*_ > *α* then model 1 is accepted
- *p_CIT_*,_*x*→*y*_ > *α* and *p*_CIT, *y*→*x*_ < *α* then model 2 is accepted
- *p_CIT_*,_*x*→*y*_ > *α* and *p*_CIT, *y*→*x*_ > *α* then no evidence for a causal relationship
- *p_CIT_*,_*x*→*y*_ < *α* and *p*_CIT, *y*→*x*_ < *α* then the confounding model is accepted (*x* ← *g* → *y*).

For the purposes of compiling simulation results we use an arbitrary *α* = 0.05 value, though we stress that for real analyses it is not good practice to rely on p-values for making causal inference, nor is it reliable to depend on arbitrary significance thresholds (37).

### MR causal test

Two stage least squares (2SLS) is a commonly used technique for performing MR when the exposure, outcome and instrument data are all available in the same sample. A p-value for this test, *p_MR_,* was obtained using the *systemfit* function in the R package *R/systemfit* (38). Note that the value of *p_MR_* is identical when using the same genetic variant to instrument the influence of the exposure *x* on the outcome *y*, or erroneously, instrumenting the outcome *y* on the exposure *x*.

The method that we will now describe is designed to distinguish between two models, *x* → *y* or *y* → *x.* Unlike the CIT framework, this approach cannot infer if the true model is *x* ← *g* → *y*. We also assume all genetic effects are additive.

To infer the direction of causality it is desirable to know which of the variables, *x* or *y*, is being directly influenced by the instrument *g.* This can be achieved by assessing which of the two variables has the biggest absolute correlation with *g* (Appendix 2), formalised by testing for a difference in the correlations *ρ_gx_* and *ρ_gy_* using Steiger’s Z-test for correlated correlations within a population (39). It is calculated as

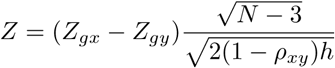

where Fisher’s z-transformation is used to obtain 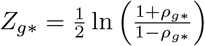,

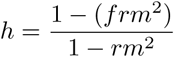

where

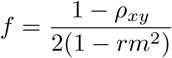

and

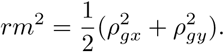

The *Z* value is interpreted such that

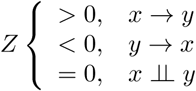

and a p-value, *p_steiger_* is generated from the *Z* value to indicate the probability of obtaining a difference between correlations *ρ_gx_* and *ρ_gy_* at least as large as the one observed, under the null hypothesis that both correlations are identical.

The existence of causality and its direction is inferred based on combining information from the MR analysis and the Steiger test. The MR analysis indicates whether there is a potential causal relationship (*p*_MR_), and the Steiger test indicates the direction (*sign*(*Z*)) of the causal relationship and the confidence of the direction (*p_steiger_*). For the purposes of compiling simulation results, these can be combined using an arbitrary *α =* 0.05 value:

- If *p_steiger_* < *α* and *p_MR_* < *α* and *Z* > 0 then a causal association for the correct model is accepted, *x* → *y*
- If *p_steiger_* < *α* and *p_MR_* < *α* and *Z* < 0 then a causal association for the incorrect model is accepted, *y* → *x*
- Otherwise if *p_steiger_* > *α* or *p_MR_* > *α,* neither model is accepted

Note that the same correlation test approach can be applied to a two-sample MR (28) setting. Two-sample MR refers to the case where the SNP-exposure association and SNP-outcome association are calculated in different samples (e.g. from publicly available summary statistics). Here the Steiger test of two independent correlations can be applied where.

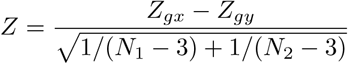

An advantage of using the Steiger test in the two sample context is that it can compare correlations in independent samples where sample sizes are different. Steiger test statistics were calculated using the *r.test* function in the R package *R/psych* (40).

### Causal direction sensitivity analysis

The Steiger test for inferring if *x* → *y* is based on evaluating *ρ _gx_ > ρ _gy_*. However, *ρ _gx_* (or *ρ _gy_*) are underestimated if *x* (or *y*) are measured imprecisely. If, for example, *x* has lower measurement precision than *y* then we might empirically obtain *ρ_g,x_o__* < *ρ_g,y_o__* because *ρ_g,x_o__* could be underestimated more than *ρ_g,y_o__*.

As we show in Appendix 2 it is possible to infer the bounds of measurement error on *x*_0_ or *y*_0_ given known genetic associations. The maximum measurement imprecision of *x*_0_ is *ρ_g,x_o__*, because it is known that at least that much of the variance has been explained in *x*_0_ by *g.* The minimum is 0, denoting perfectly measured trait values (the same logic applies to *y*_0_). It is possible to simulate what the inferred causal direction would be for all values within these bounds.

To evaluate how reliable, *R,* the inference of the causal direction is to potential measurement error in *x* and *y* we need to predict the values of *ρ_gy_* − *ρ_gx_* for those values of measurement error. We integrate over the entire range of *ρ_gy_* − *ρ_gx_* values for possible measurement error values. We find the ratio of the volume that agrees with the inferred direction of causality over the volume that disagrees with the inferred direction of causality. A ratio *R* = 1 indicates that the inferred causal direction is highly sensitive to measurement error, with equal weight of the measurement error parameter space supporting both directions of causality. In general, the *R* value denotes that the inferred direction of causality is *R* times more likely to be the empirical result than the opposite direction. Full details are provided in Appendix 2.

### Simulations

Simulations were conducted by creating variables of sample size *n* for the exposure *x*, the measured values of the exposure *x*_o_, the outcome *y*, the measured values of the outcome *y*_o_ and the instrument *g.* One of two models are simulated, the “causal model” where *x* causes *y* and *g* is an instrument for x; or the “non-causal model” where *g* influences a confounder *u* which in turn causes both *x* and *y*. Here *x* and *y* are correlated but not causally related. Each variable in the causal model was simulated such that:

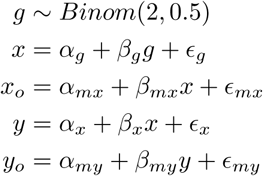

where 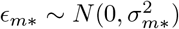, *α_mx_* and *β_mx_* are parameters that represent non-differential measurement error into the exposure variable *x*, and *α_my_* and *β_my_* are parameters for non-differential measurement error in the outcome *y*. Similarly in the non-causal model:

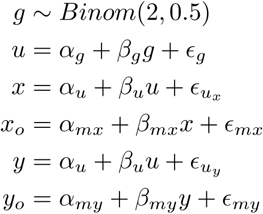

All *a* values were set to 0, and *β* values set to 1. Normally distribted values of *ϵ*_∗_ were generated such that

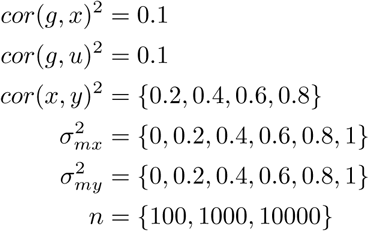

giving a total of 432 combinations of parameters. Simulations using each of these sets of variables were performed 100 times, and the CIT and MR methods were applied to each in order to evaluate the causal association of the simulated variables. Similar patterns of results were obtained for different values of *cor*(*g, x*) and *cor*(*g, u*).

### Applied example using two sample MR

Two sample MR (28) was performed using the summary statistics for genetic influences on gene expression and DNA methylation. To do this we obtained a list of 458 gene expression - DNA methylation associations as reported in Shakhbazov et al (41). These were filtered to be located on the same chromosome, have robust correlations after correcting for multiple testing, and to share a SNP that had a robust cis-acting effect on both the DNA methylation probe and the gene expression probe. Because only summary statistics were available (effect, standard error, effect allele, sample size, p-values) for the instrumental SNP on the methylation and gene expression levels, the Steiger test of two independent correlations was used to infer the direction of causality for each of the associations. The Wald ratio test was then used to estimate the causal effect size for the estimated direction for each association.

All analysis was performed using the R programming language (42) and code is made available at https://github.com/explodecomputer/causal-directions and implemented in the MR-Base (http://wwww.mrbase.org) platform (26).

## Results

### Mediation-based causal inference under measurement error

In the causal inference test (CIT), the 4th condition (see Methods) employs mediation for causal inference, and can be expressed as *cov*(*g, y* − *ŷ*) = 0, where *ŷ* = *α ̂_x_ + β̂_x_x*_o_. When measurement error in scale and imprecision is introduced, such that *y*_o_ is the measured value of *y*, it can be shown using basic covariance properties (Appendix 1) that

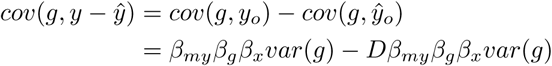

where

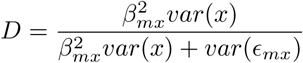

Thus an observational study will find *cov*(*g, y_o_* – *ŷ_o_*) = 0 when the true model is causal only when *D* = 1. Therefore, if there is any measurement error that incurs imprecision in *x* (i.e. *var*(*ϵ_mx_*) ≠ 0) then there will remain an association between *g* and *y_o_*|*x_o_*, which is in violation of the the 4th condition of the CIT. Note that scale transformation of *x* or *y* without any incurred imprecision is insufficient to lead to a violation of the test statistic assumptions, and henceforth mention of measurement error will relate to imprecision unless otherwise stated.

We performed simulations to verify that this problem does arise using the CIT method. Figure 2 shows that when there is no measurement error in the exposure or outcome variables (*ρ_x,x_o__* = 1) the CIT is reliable in identifying the correct causal direction. However, as measurement error increases in the exposure variable, eventually the CIT is more likely to infer a robust causal association in the wrong direction. Also of concern here is that increasing sample size does not solve the issue, indeed it only strengthens the apparent evidence for the incorrect inference.

**Figure 2.**
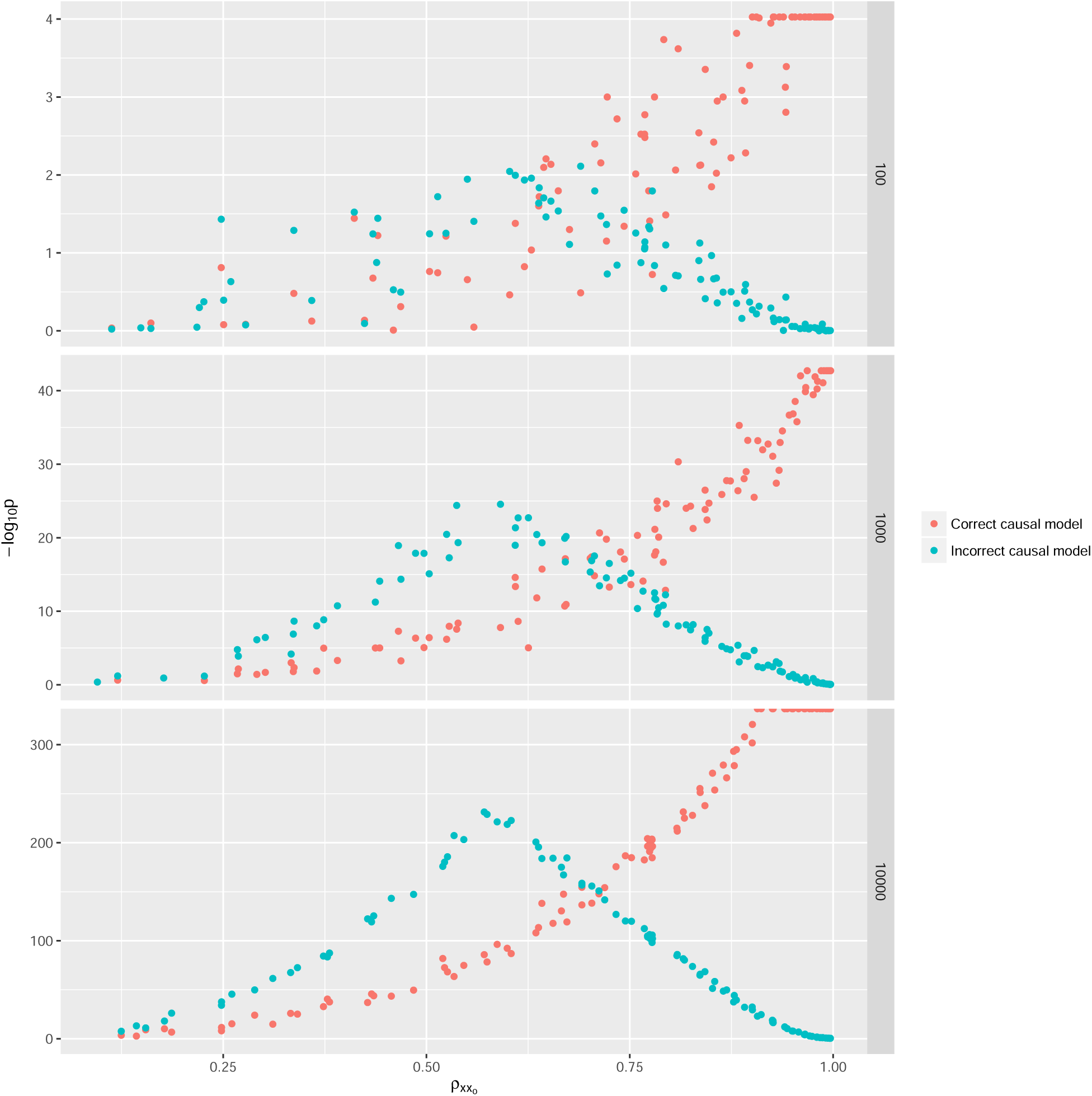
The CIT was performed on simulated variables where the exposure influenced the outcome and the exposure was instrumented by a SNP. The test statistic from CIT when testing if the exposure caused the outcome (the true model) is in red, and the test for the outcome causing the exposure (false model) is in green. Rows of plots represent the sample sizes used for the simulations. As measurement imprecision increases (decreasing values on x-axis) the test statistic for the incorrect model gets stronger and the test statistic for the correct model gets weaker.

We also performed simulations to compare the performance of MR against CIT in detecting a causal association between simulated variables under different levels of imprecision simulated in the exposure. Figure 3 shows the true positive rates between the CIT and MR for detecting a causal association. We observe that the CIT has lower power in all cases, with performance declining as measurement imprecision increases in the exposure. When MR assumptions are satisfied, notably that it is known on which of *x* and *y* the SNP *g* has a primary influence, the performance of MR in detecting an association is unrelated to measurement error in the exposure. Measurement error in the outcome does reduce power, but does not induce a substantive difference in performance between CIT and MR.

**Figure 3.**
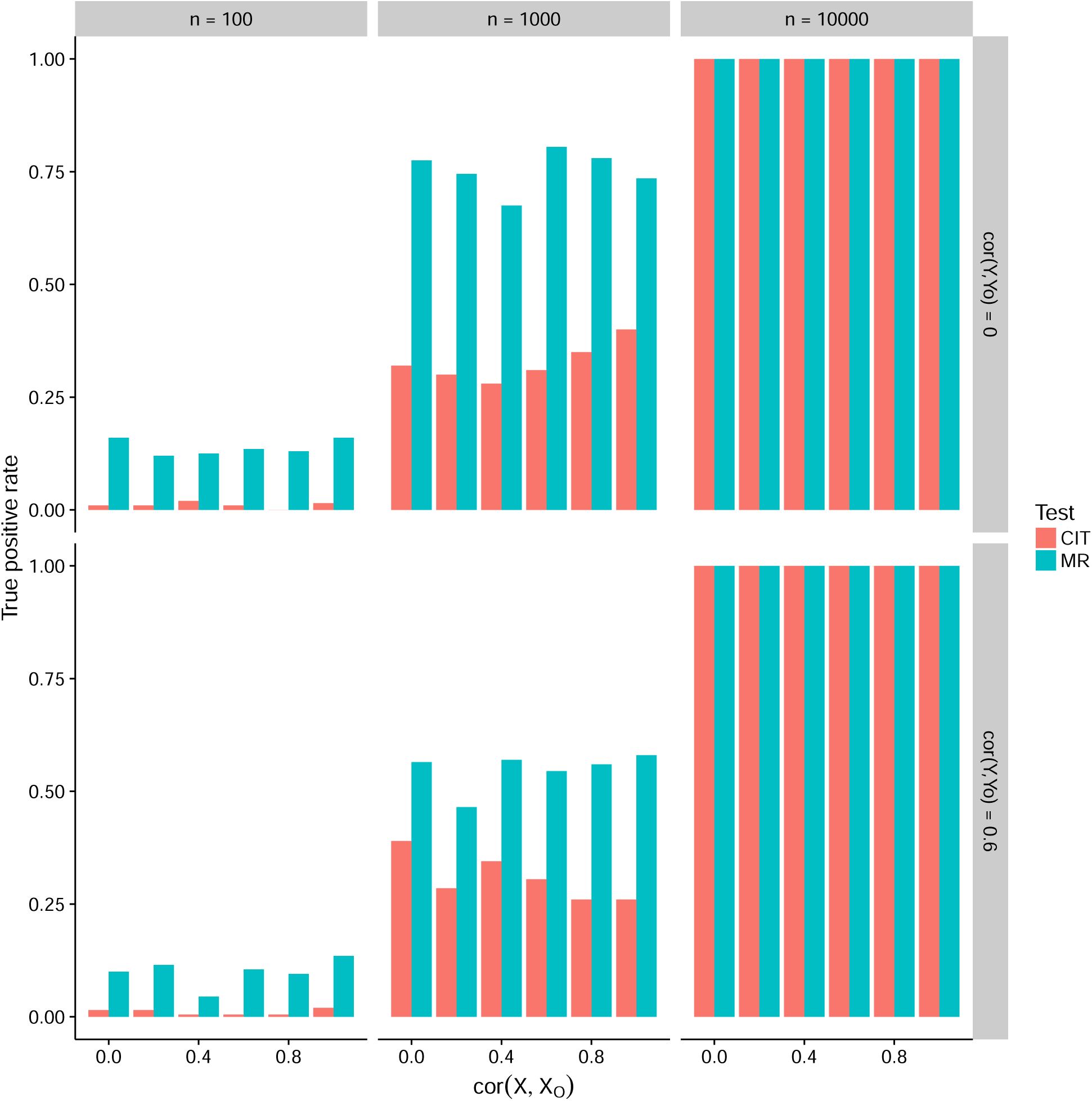
Outcomes were simulated to be causally influenced by exposures with varying degrees of measurement imprecision applied to the exposure variable (x axis). True positive rates (y axis) for MR and CIT were compared for varying levels of measurement imprecision in the outcome variable (rows of boxes) and sample sizes (columns of boxes).

### Using MR Steiger to infer the direction of causality

If we do not know whether the SNP *g* has a primary influence on *x* or *y* then CIT can attempt to infer the causal direction. Here we introduce the MR Steiger approach to similarly orient the direction of causality but in an MR analysis when the underlying biology of the SNP is not fully understood.

For a particular association, it is of interest to identify the range of possible measurement error values agree and disagree with the empirically inferred causal direction (Figure 4a, Appendix 2). This metric can be used to evaluate the reliability of MR Steiger.

**Figure 4.**
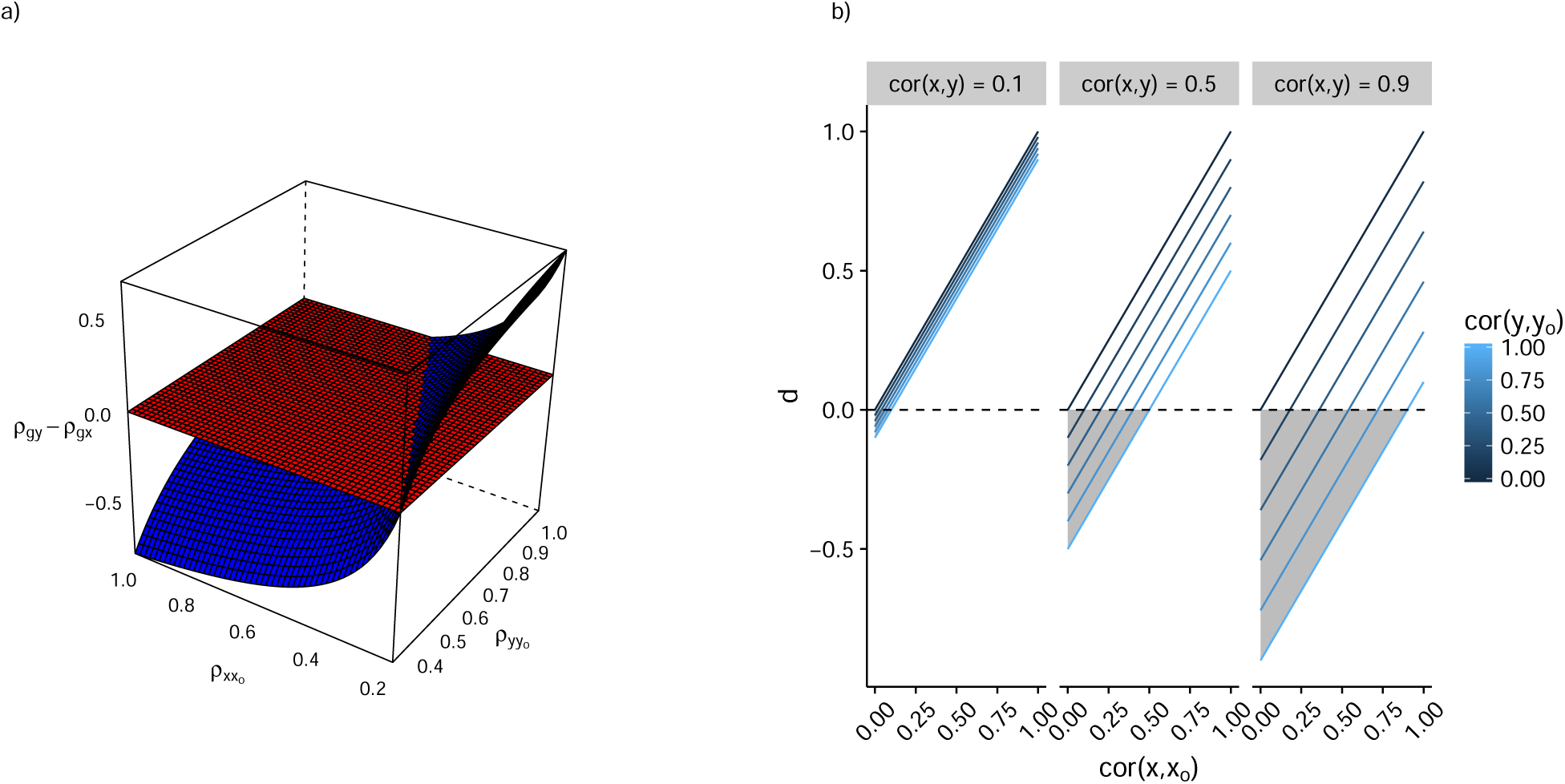
a) We can predict the values the Steiger test would take (z-axis) for different potential values of measurement error (x and y axes), drawn here as the blue surface. When *ρ _g y_ > ρ _g_,_x_,* as denoted by the range of values where the blue surface is above the red plane, those values of measurement error lead to our observed Steiger test inferring the wrong causal direction. Where the blue surface lies below the red plane, these measurement error values support the inferred causal direction of X to Y. A measure of reliability, therefore, is the ratio of the negative and positive volumes of the total space bound by the blue and red surfaces, 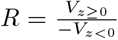. In this case, where 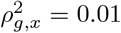 and 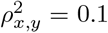, the *R* = 4.40, which means that 4.40 times as much of the possible measurement error values are in support of the *x* → *y* direction of causality than *y* → *x.* b) Plots depicting the parameter space in which the function *d* = *cor*(*x, x_O_*) − *cor*(*x, y*)*cor*(*y, y_O_*) is negative. When *d* is negative the Steiger test is liable to infer the wrong direction of causality. Shaded regions show the parameter space where *d* is negative. The graph shows that for the majority of the parameter space of the function, *d* is positive, especially where causal relationships are relatively weak.

We show that in the presence of measurement imprecision, *d* = *ρ_x,y_ρ_y,y_o__* (Appendix 2) determines the range of parameters around which the Steiger test is liable to provide the wrong direction of causality *(i.e.* if *d* > 0 then the Steiger test is likely to be correct about the causal direction). Figure 4b shows that when there is no measurement error in *x*, the Steiger test is unlikely to infer the wrong direction of causality even if there is measurement error in *y.* It also shows that in most cases where *x* is measured with error, especially when the causal effect between *x* and *y* is not very large, the sensitivity of the Steiger test to measurement error is relatively low.

### Comparison of CIT and MR Steiger for obtaining the correct direction of causality

We used simulations to explore the performance of the MR Steiger approach in comparison to CIT in terms of the rate at which evidence of a causal relationship is obtained for the correct direction of causality, and the rate at which evidence of a causal relationship is obtained where the reported direction of causality is incorrect. Simulations were performed for two models, one for a “causal model” where there was a causal relationship between *x* and y; and one for a “non-causal model” where *x* and *y* were not causally related, but had a confounded association induced by the SNP *g* influencing a confounder variable *u*.

Figure 5a shows that, for the “causal model”, the MR analysis is indeed liable to infer the wrong direction of causality when *d* < 0, and that this erroneous result is more likely to occur with increasing sample size. However, the CIT is in general more fallable to reporting a robust causal association for the wrong direction of causality. When *d* > 0 we find that in most cases the MR method has greater power to obtain evidence for causality than CIT, and always obtains the correct direction of causality. The CIT, unlike the Steiger test for MR, is able to distinguish the “non-causal model” from the “causal model” (Methods, Figure 5b), but it is evident that measurement error will often lead the CIT to identify the causal model as true, when in fact the underlying model is this non-causal model.

**Figure 5.**
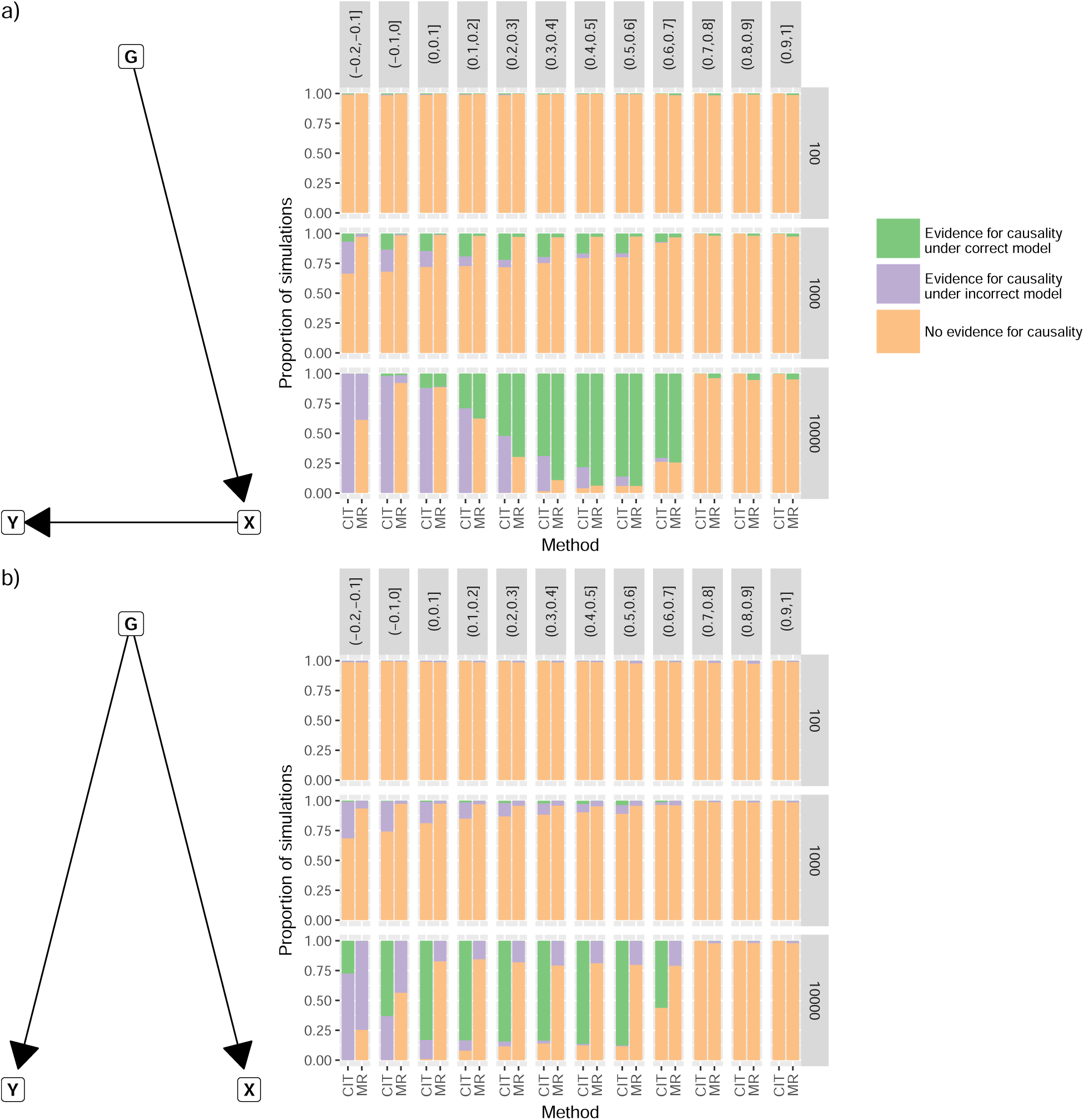
a) Outcome *y* was simulated to be caused by exposure *x* as shown in the graph, with varying degrees of measurement error applied to both. CIT and MR were used to infer evidence for causality between the exposure and outcome, and to infer the direction of causality. The value of *d* = *ρ_x,x_o__* − *ρ_x,y_ρ_y,y_o__*, such that when *d* is negative we expect the Steiger test to be more likely to be wrong about the direction of causality. Rows of graphs represent the sample size used in the simulations. For the CIT method, outcome 1 denoted evidence for causality with correct model, outcomes 2 or 3 denoted evidence for causality with incorrect model, and outcome 4 denoted no evidence for causality. b) As in (a) except the simulated model was non-causal, and a genetic confounder induces an association between *x* and *y*. MR is unable to identify this model, so any significant associations are deemed to be incorrect. Outcome 3 denotes evidence for the correct model for the CIT method.

### The causal relationship between gene expression and DNA methylation levels

We used the Steiger test to infer the direction of causality between DNA methylation and gene expression levels between 458 putative associations. We found that the causal direction commonly goes in both directions (Figure 6a), but assuming no or equal measurement error, DNA methylation levels were the predominant causal factor (*p =* 1.3 × 10^−5^). The median reliability (R) of the 458 tests was 3.92 (5%-95% quantiles 1.08 - 37.11). We then went on to predict the causal directions of the associations for varying levels of systematic measurement error for the different platforms. Figure 6a shows that the conclusions about the direction of causality between DNA methylation and gene expression are very sensitive to measurement error.

**Figure 6.**
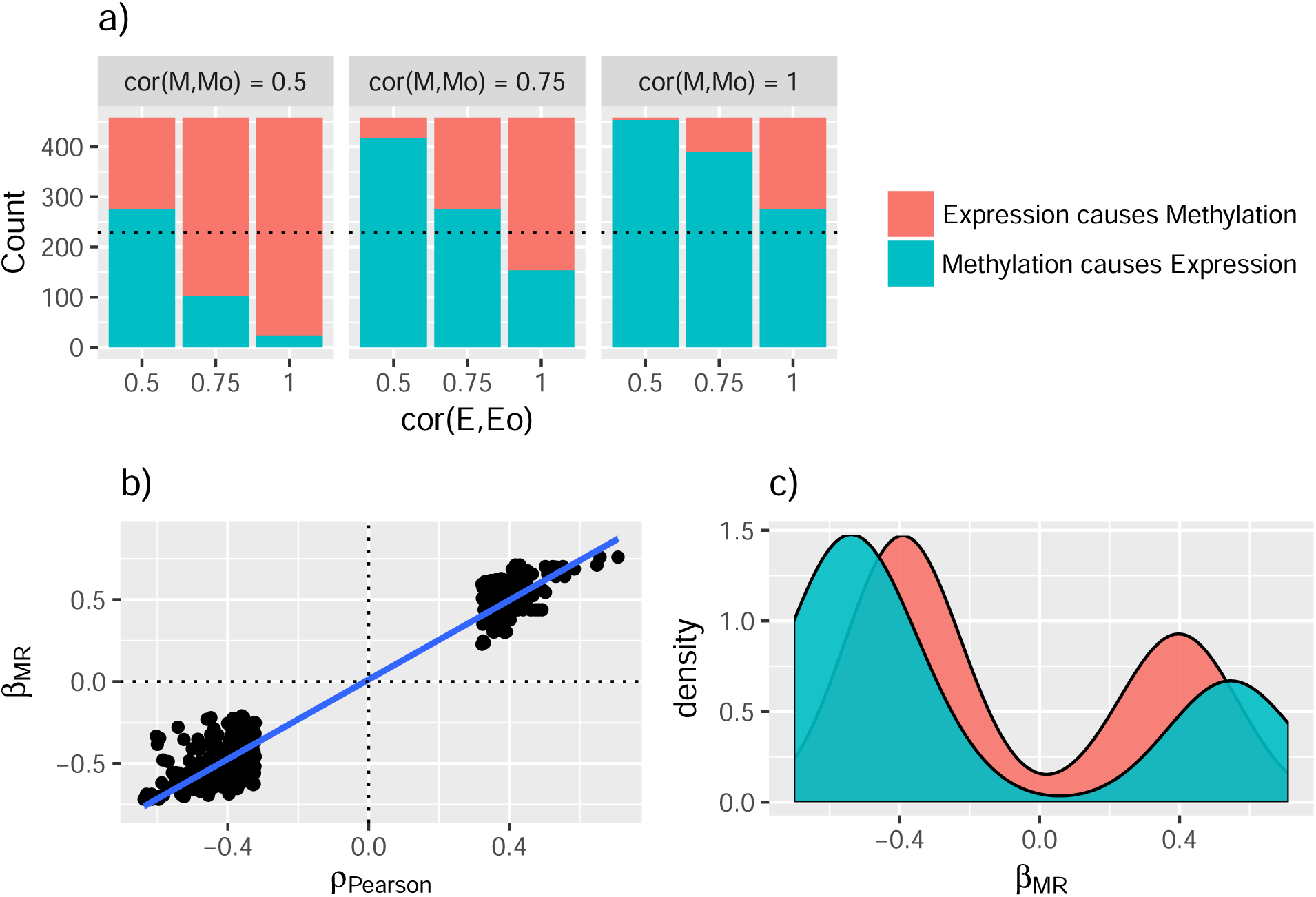
Using 458 putative associations between DNA methylation and gene expression we used the Steiger test to infer the direction of causality between them. a) The rightmost bar shows the proportion of associations for each of the two possible causal directions (colour key) assuming no measurement error in either gene expression or DNA methylation levels. The proportions change when we assume different levels of measurement error in gene expression levels (x-axis) or DNA methylation levels (columns of boxes). If there is systematically higher measurement error in one platform than the other it will appear to be less likely to be the causal factor. b) The relationship between the Pearson correlation between DNA methylation and gene expression levels (x-axis) and the causal estimate (scaled to be in standard deviation units, y-axis). c) Distribution of estimated causal effect sizes, stratified into associations inferred to be due to DNA methylation causing expression (blue) and expression causing DNA methylation (red).

We performed two sample MR (28) for each association in the direction of causality inferred by the Stieger test. We observed that the sign of the MR estimate was generally in the same direction as the Pearson correlation coefficient reported by Shakhbazov et al (41) (Figure 6b). There was a moderate correlation between the absolute magnitudes of the causal correlation and the observational Pearson correlation (r = 0.45). Together these inferences suggest that even in estimating associations between ‘omic’ variables, which are considered to be low level phenotypes, it is important to use causal inference methods over observational associations to infer causal effect sizes.

We also observed that for associations where methylation caused gene expression the causal effect was more likely to be negative than for the associations where gene expression caused methylation (OR = 0.61 (95% CI 0.36 - 1.03), Figure 6c), suggesting that reducing methylation levels at a controlling CpG typically leads to increased gene expression levels, consistent with expectation (43).

## Discussion

Researchers are often confronted with the problem of making causal inferences using a statistical framework on observational data. In the epidemiological literature issues of measurement error in mediation analysis are relatively well explored (44). Our analysis extends this to related methods such as CIT that are widely used in predominantly ’omic data. These methods are indeed susceptible to the same problem as standard mediation based analysis, and specifically we show that as measurement error in the (true) exposure variable increases, CIT is likely to have reduced statistical power, and liable to infer the wrong direction of causality.

We also demonstrate that, though unintuitive, increasing sample size does not resolve the issue, rather it leads to more extreme p-values for the model that predicts the wrong direction of causality.

Under many circumstances a practical solution to this problem is to use Mendelian randomisation instead of methods such as the CIT or similar that are based on mediation. Inferring the existence of causality using Mendelian randomisation is robust in the face of measurement error and, if the researcher has knowledge about the biology of the instrument being used in the analysis, can offer a direct solution to the issues that the CIT faces. This assumption is often reasonable, for example SNPs are commonly used as instruments when they are found in genes with known biological relevance for the trait of interest. But on many occasions, especially in the realm of ’omic data, this is not the case, and methods based on mediation have been valuable in order to be able to both ascertain if there is a causal association and to infer the direction of causality. Here we have described a simple extension to MR which can be used as an alternative to or in conjunction with mediation based methods. We show that this method is still liable to measurement error, but because it has different properties to the CIT it offers several main advantages. First, it uses a formal statistical framework to test for the reliability of the assumed direction of causality. Second, after testing in a comprehensive range of scenarios the MR based approach is less likely to infer the wrong direction of causality compared to CIT, while substantially improving power over CIT in the cases where *d* > 0.

We demonstrate this new method by evaluating the causal relationships of 458 known associations between DNA methylation and gene expression levels using summary level data. The inferred causal direction is heavily influenced by how much measurement error is present in the different assaying platforms. For example, if DNA methylation measures typically have higher measuremet error than gene expression measures then our analysis suggests that DNA methylation levels would be more often the causal factor in the association. Indeed, previous studies which have evaluated measurement error in these platforms do support this position (45,46), though making strong conclusions for this analysis is difficult because measurement error is likely to be study specific. We also haven’t accounted for the influence of winner’s curse, which can inflate estimates of the variance explained by SNPs, with higher inflation expected amongst lower powered studies. Using p-values for genetic associations from replication studies will mitigate this problem.

In our simulations we focused on the simple case of a single instrument in a single sample setting with a view to making a fair comparison between MR and the various mediation-based methods available. However, if there is only a single instrument it is difficult to separate between the two competing models of *g* instrumenting a trait which causes another trait, and *g* having pleiotropic effects on both traits independently (47). Under certain conditions of measurement error the CIT test can distinguish these models. We also note that it is straightforward to extend the MR Steiger approach to multiple instruments, requiring only that the total variance explained by all instruments be calculated under the assumption that they are independent. Multiple instruments can indeed help to distinguish between the causal and pleiotropic models, for example by evaluating the proportionality of the SNP-exposure and SNP-outcome effects (16). Additionally, if there is at least one instrument for each trait then bi-directional MR can offer solutions to inferring the causal direction (16,48,49). We restricted the simulations to evaluating the causal inference between quantitative traits, but it is possible that the analysis could be extended to binary traits by using the genetic variance explained on the liability scale, taking into account the population prevalence (50). However, our analysis goes beyond many previous explorations of measurement error by assessing the impacts of both imprecision (noise) and linear transformations of the true variable on causal inference.

In this work we assumed that pleiotropy (the influence of the instrument on the outcome through a mechanism other than the exposure) was not present. Recent method developments in MR (24,25) have focused on accounting for the issues that horizontal pleiotropy can introduce when multiple instruments are available, but how they perform in the presence of measurement error remains to be explored. An important advantage that MR confers over most mediation based analysis is that it can be performed in two samples, which can considerably improve power and expand the scope of analysis. However, whether there is a substantive difference in two sample MR versus one sample MR in how measurement error has an effect is not yet fully understood. We have also assumed no measurement error in the genetic instrument, which is not unreasonable given the strict QC protocols that ensure high quality genotype data is available to most studies. We have restricted the scope to only exploring non-differential measurement error and avoided the complications incurred if measurement error in the exposure and outcome is correlated. We have also not addressed other issues pertaining to instrumental variables which are relevant to the question of instrument-exposure specification. One such problem is exposure misspecification, for example an instrument could associate with several closely related putative outcomes, with only one of them actually having a causal effect on the outcome. This problem has shown to be the case for SNPs influencing different lipid fractions, for example (51,52).

Mediation based network approaches, that go beyond analyses of two variables, are very well established (35) and have a number of extensions that make them valuable tools, including for example network construction. But because they are predicated on the basic underlying principles of mediation they are liable to suffer from the same issues of measurement error. Recent advances in MR methodology, for example applying MR to genetical genomics (53), multivariate MR (52) and mediation through MR (54-56) may offer more robust alternatives for these more complicated problems.

The overarching result from our simulations is that, regardless of the method used, inferring the causal direction using an instrument of unknown biology is highly sensitive to measurement error. With the presence of measurement error near ubiquitous in most observational data, and our ability to measure it limited, we argue that it needs to be central to any consideration of approaches which are used in attempt to strengthen causal inference, and any putative results should be accompanied with appropriate sensitivity analysis that assesses their robustness under varying levels of measurement error.

## Acknowledgements

This work was funded by the UK Medical Research Council (MC_UU_12013/1 and MC_UU_12013/9).

## Appendix 1

We assume the following model

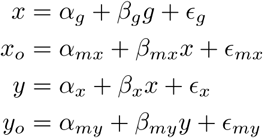

where *x* is the exposure on the outcome *y*, *g* is an instrument that has a direct effect on *x*, *x*_0_ is the measured quantity of *x*, where measurement error is incurred from linear transformation in *α_mx_* and *β_mx_* and imprecision from *ϵ _mx_*, and *y_o_* is the measured quantity of *y*, where measurement error is incurred from linear transformation in *α_my_* and *β_my_* and imprecision from *ϵ_my_*. Our objective is to estimate the expected magnitude of association between *g* and *y* after conditioning on *x*. Under the CIT, this is expected to be *cov*(*g, y_o_* − *ŷ_o_*) = 0 when *x* causes *y*, where *ŷ_o_* = *â_x_o__* + *β̂_x_o__x_o_* is the predicted value of *y_o_* using the measured value of *x_o_.*

We can split *cov*(*g, y_o_* − *ŷ_o_*) into two parts, *cov*(*g, y_o_*) and *cov*(*g, ŷ*_o_).

### Part 1

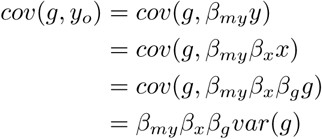

### Part 2

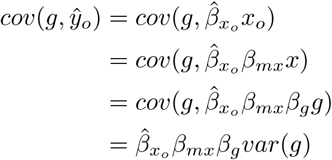

Simpifying further

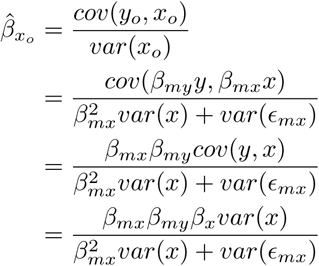

which can be substituted back to give

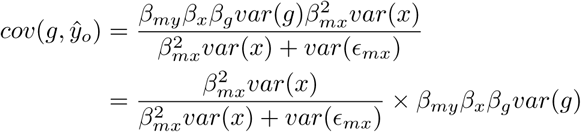

Finally

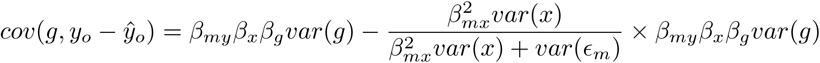

thus *cov*(*g, y_o_* − *ŷ_o_*) = 0 if the measurement imprecision in *x_o_* is *var*(*ϵ_m_*) = 0. However if there is any imprecision then the condition *cov*(*g, y_o_ − *ŷ*_o_*) = 0 will not hold.

## Appendix 2

Assuming that either *x* → *y* or *y* → *x*, the causal direction can be inferred by evaluating which of *ρ_g_,_x_* and *ρ_g,y_* is larger in magnitude. The Steiger test is a hypothesis test that provides a p-value for observing the difference in these correlations under the null hypothesis that they are equal.

Assuming the causal direction is *x* → *y*, two stage MR is formulated using the following regression models:

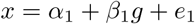

for the first stage and

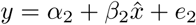

where *x ̂* = *α̂_1_ + β ̂*_1_*g.* Writing in scale free terms, *ρ_g x_* denotes the correlation between *g* and the exposure variable *x*, and it is expected that *ρ_g_,_x_ > ρ_g_,_y_* because *ρ_g_,_y_* = *ρ_g_,_x_ρ_x_,_y_*, where *ρ_x_,_y_* is the causal association between *x* and *y* (which is likely to be less than 1).

In the presence of measurement error in *x* and *y*, however, the empirical inference of the causal direction will instead be based on evaluating *ρ_g,x_o__* > *ρ_g,y_o__*, which can be simplified:

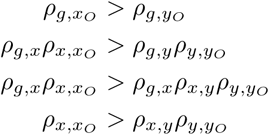

In order to assess how reliable the inference of the causal direction is in the presence of measurement imprecision, we can evaluate the range of potential values of measurement error in *x* and *y* over which the empirical difference in *ρ_g,x_o__* and *ρ_g,y_o__* would return the wrong causal direction.

For different values of *ρ_x,x_o__*, 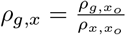 and *ρ_g,x_o__* ≤ *ρ_x,x_o__* ≤ 1. For different values of *ρ_y,y_o__*, 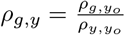 and *ρ_g,y_o__* ≤ *ρ_y,x_o__* ≤ 1

Call *z* = *ρ_g y_* − *ρ_g x_* the true difference in the variance explained by the genetic variant in *y* and *x*. If *z* < 0 then we infer that *x* → *y*. There will be some values of *ρ_x,x_o__* and *ρ_y,y_o__* that do not alter whether *z* < 0. To evaluate the reliability, *R,* of the inference of the causal direction with regards to measurement error, the objective is to compare the proportion of the parameter space that agrees with the inferred direction against the proportion which does not:

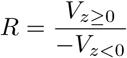

If *R* =1 then the direction of causality is equally probable across the range of possible measurement error values. If *R* > 1 then *R* times as much of the parameter space favours the inferred direction of causality. *V*_z_, the total volume of the function (Figure 4), can be obtained analytically by solving:

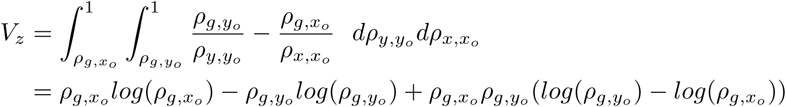

V_*z* ≥ 0_, the proportion of the volume that lies above the *z* = 0 plane, can also be obtained analytically. The region of this volume is bound by the values of *ρ_x,x_o__* and *ρ_y,y_o__* where 0 = *ρ_g,y_* − *ρ_g,x_*, which can be expanded to *ρ_y,y_o__* = *ρ_g,y_o__ρ_x,x_o__*/*ρ_g,x_o__*. Hence,

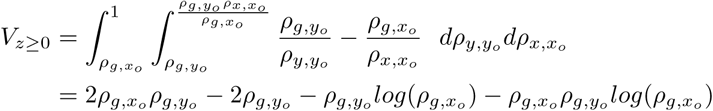

Thus *V*_*z* < 0_ = *V_z_* — *V*_*z* ≥ 0_.

## References

1. Phillips AN, Davey Smith G. How independent are “independent” effects? relative risk estimation when correlated exposures are measured imprecisely. Journal of Clinical Epidemiology. 1991 44(11):1223–31.

2. Davey Smith G, Ebrahim S. Data dredging, bias, or confounding. BMJ. 2002 325(7378):1437–8.

3. Davey Smith G, Ebrahim S. Mendelian randomization: prospects, potentials, and limitations. International journal of epidemiology [Internet]. 2004 Feb;33(1):30–42. Available from: http://www.ncbi.nlm.nih.gov/pubmed/15075143

4. Millstein J, Zhang B, Zhu J, Schadt EE. Disentangling molecular relationships with a causal inference test. BMC genetics [Internet]. 2009 Jan;10:23. Available from: http://www.pubmedcentral.nih.gov/articlerender.fcgi?artid=3224661{\&}tool=pmcentrez{\&}rendertype=abstract

5. Aten JE, Fuller TF, Lusis AJ, Horvath S. Using genetic markers to orient the edges in quantitative trait networks: the NEO software. BMC systems biology [Internet]. 2008 Jan;2:34. Available from: http://www.pubmedcentral.nih.gov/articlerender.fcgi?artid=2387136{\&}tool=pmcentrez{\&}rendertype=abstract

6. Waszak SM, Delaneau O, Gschwind AR, Kilpinen H, Raghav SK, Witwicki RM, et al. Variation and genetic control of chromatin architecture in humans. Cell [Internet]. 2015 162(5):1039–50. Available from: http://dx.doi.org/10.1016/j.cell.2015.08.001

7. Houle D, Pélabon C, Wagner G, Hansen T. Measurement and meaning in biology. The Quarterly Review of Biology [Internet]. 2011 86(1):3–34. Available from: http://www.jstor.org/stable/10.1086/658408

8. Hernán M a, Cole SR. Invited Commentary: Causal diagrams and measurement bias. American journal of epidemiology [Internet]. 2009 Oct;170(8):959–62; discussion 963–4. Available from: http://www.pubmedcentral.nih.gov/articlerender.fcgi?artid=2765368{\&}tool=pmcentrez{\&}rendertype=abstract

9. Harper KN, Peters B a, Gamble MV. Batch effects and pathway analysis: two potential perils in cancer studies involving DNA methylation array analysis. Cancer epidemiology, biomarkers & prevention: a publication of the American Association for Cancer Research, cosponsored by the American Society of Preventive Oncology [Internet]. 2013 Jun;22(6):1052–60. Available from: http://www.pubmedcentral.nih.gov/articlerender.fcgi?artid=3687782{\&}tool=pmcentrez{\&}rendertype=abstract

10. Chen Y-a, Lemire M, Choufani S, Butcher DT, Grafodatskaya D, Zanke BW, et al. Discovery of cross-reactive probes and polymorphic CpGs in the Illumina Infinium HumanMethylation450 microarray. Epigenetics: official journal of the DNA Methylation Society [Internet]. 2013 Feb;8(2):203–9. Available from: http://www.pubmedcentral.nih.gov/articlerender.fcgi?artid=3592906{\&}tool=pmcentrez{\&}rendertype=abstract

11. Houseman EA, Accomando WP, Koestler DC, Christensen BC, Marsit CJ, Nelson HH, et al. DNA methylation arrays as surrogate measures of cell mixture distribution. BMC bioinformatics [Internet]. 2012 Jan;13:86. Available from: http://www.pubmedcentral.nih.gov/articlerender.fcgi?artid=3532182{\&}tool=pmcentrez{\&}rendertype=abstract

12. Ahima RS, Lazar MA. Physiology. The health risk of obesity-better metrics imperative. Science (New York, NY) [Internet]. 2013 Aug;341(6148):856–8. Available from: http://science.sciencemag.org/content/341/6148/856.abstract

13. Cessie S le, Debeij J, Rosendaal FR, Cannegieter SC, Vandenbroucke JP. Quantification of bias in direct effects estimates due to different types of measurement error in the mediator. Epidemiology (Cambridge, Mass) [Internet]. 2012 Jul;23(4):551–60. Available from: http://www.ncbi.nlm.nih.gov/pubmed/22526092

14. Blakely T, McKenzie S, Carter K. Misclassification of the mediator matters when estimating indirect effects. Journal of epidemiology and community health [Internet]. 2013 May;67(5):458–66. Available from: http://www.ncbi.nlm.nih.gov/pubmed/23386673

15. Davey Smith G, Ebrahim S. ’Mendelian randomization’: can genetic epidemiology contribute to understanding environmental determinants of disease? International Journal of Epidemiology [Internet]. 2003 Feb;32(1):1–22. Available from: http://www.ije.oxfordjournals.org/cgi/doi/10.1093/ije/dyg070

16. Davey Smith G, Hemani G. Mendelian randomization: genetic anchors for causal inference in epidemiological studies. Human molecular genetics. 2014 Jul;23(R1):R89—R98.

17. Schadt EE, Lamb J, Yang X, Zhu J, Edwards S, GuhaThakurta D, et al. An integrative genomics approach to infer causal associations between gene expression and disease. Nature Genetics [Internet]. 2005 Jul;37(7):710–7. Available from: http://dx.doi.org/10.1038/ng1589 http://www.nature.com/ng/journal/v37/n7/full/ng1589.html http://www.nature.com/ng/journal/v37/n7/pdf/ng1589.pdf

18. Millstein J. cit: Causal Inference Test. R package version 1.9 [Internet]. 2016. Available from: http://cran.r-project.org/package=cit

19. Koestler DC, Chalise P, Cicek MS, Cunningham JM, Armasu S, Larson MC, et al. Integrative genomic analysis identifies epigenetic marks that mediate genetic risk for epithelial ovarian cancer. BMC medical genomics [Internet]. 2014 Jan;7(1):8. Available from: http://www.pubmedcentral.nih.gov/articlerender.fcgi?artid=3916313{\&}tool=pmcentrez{\&}rendertype=abstract

20. Liu Y, Aryee MJ, Padyukov L, Fallin MD, Hesselberg E, Runarsson A, et al. Epigenome-wide association data implicate DNA methylation as an intermediary of genetic risk in rheumatoid arthritis. Nature biotechnology [Internet]. 2013 Feb;31(2):142–7. Available from: http://www.pubmedcentral.nih.gov/articlerender.fcgi?artid=3598632{\&}tool=pmcentrez{\&}rendertype=abstract

21. Yuan W, Xia Y, Bell CG, Yet I, Ferreira T, Ward KJ, et al. An integrated epigenomic analysis for type 2 diabetes susceptibility loci in monozygotic twins. Nature communications [Internet]. 2014 Jan;5:5719. Available from: http://www.pubmedcentral.nih.gov/articlerender.fcgi?artid=4284644{\&}tool=pmcentrez{\&}rendertype=abstract

22. Tang Y, Axelsson AS, Spégel P, Andersson LE, Mulder H, Groop LC, et al. Genotype-based treatment of type 2 diabetes with an α2A-adrenergic receptor antagonist. Science translational medicine [Internet]. 2014 Oct;6(257):257ra139. Available from: http://www.ncbi.nlm.nih.gov/pubmed/25298321

23. Hong X, Hao K, Ladd-Acosta C, Hansen KD, Tsai H-J, Liu X, et al. Genome-wide association study identifies peanut allergy-specific loci and evidence of epigenetic mediation in US children. Nature communications [Internet]. 2015 Jan;6:6304. Available from: http://www.pubmedcentral.nih.gov/articlerender.fcgi?artid=4340086{\&}tool=pmcentrez{\&}rendertype=abstract

24. Bowden J, Davey Smith G, Burgess S. Mendelian randomization with invalid instruments: effect estimation and bias detection through Egger regression. International Journal of Epidemiology. 2015 44(2):512–25.

25. Bowden J, Davey Smith G, Haycock PC, Burgess S. Consistent Estimation in Mendelian Randomization with Some Invalid Instruments Using a Weighted Median Estimator. Genetic Epidemiology [Internet]. 2016 May;40(4):304–14. Available from: http://www.ncbi.nlm.nih.gov/pubmed/27061298 http://www.pubmedcentral.nih.gov/articlerender.fcgi?artid=PMC4849733 http://doi.wiley.com/10.1002/gepi.21965

26. Hemani G, Zheng J, Wade KH, Laurin C, Elsworth B, Burgess S, et al. MR-Base: a platform for systematic causal inference across the phenome using billions of genetic associations. BioRxiv. 2016;10.1101/07.

27. Claussnitzer M, Dankel SN, Kim K-H, Quon G, Meuleman W, Haugen C, et al. FTO Obesity Variant Circuitry and Adipocyte Browning in Humans. The New England journal of medicine. 2015 373(10):895–907.

28. Pierce BL, Burgess S. Efficient design for Mendelian randomization studies: subsample and 2-sample instrumental variable estimators. American journal of epidemiology [Internet]. 2013 Oct;178(7):1177–84. Available from: http://www.pubmedcentral.nih.gov/articlerender.fcgi?artid=3783091{\&}tool=pmcentrez{\&}rendertype=abstract

29. Hernán MA, Hernández-Díaz S, Robins JM. A structural approach to selection bias. Epidemiology (Cambridge, Mass) [Internet]. 2004 Sep;15(5):615–25. Available from: http://www.ncbi.nlm.nih.gov/pubmed/15308962

30. Pierce BL, VanderWeele TJ. The effect of non-differential measurement error on bias, precision and power in Mendelian randomization studies. International Journal of Epidemiology [Internet]. 2012 Oct;41(5):1383–93. Available from: https://academic.oup.com/ije/article-lookup/doi/10.1093/ije/dys141

31. Ashenfelter O, Krueger AB. Estimates of the Economic Return to Schooling from a New Sample of Twins. The American Economic Review. 1994 84(5):1157–73.

32. Nagarajan R, Scutari M. Impact of noise on molecular network inference. PloS one [Internet]. 2013 Jan;8(12):e80735. Available from: http://www.pubmedcentral.nih.gov/articlerender.fcgi?artid=3855153{\&}tool=pmcentrez{\&}rendertype=abstract

33. Shpitser I, VanderWeele T, Robins J. On the validity of covariate adjustment for estimating causal effects. Proceedings of the Twenty Sixth Conference on Uncertainty in Artificial Intelligence (UAI-10). 2010;527–36.

34. Wang L, Michoel T. Detection of regulator genes and eQTLs in gene networks. arXiv [Internet]. 2015 Dec;arXiv:1512. Available from: http://arxiv.org/abs/1512.05574

35. Lagani V, Triantafillou S, Ball G, Tegner J, Tsamardinos I. Probabilistic Computational Causal Discovery for Systems Biology. In: Uncertainty in biology: A computational modeling approach [Internet]. Springer; 2015. p. 47. Available from: https://books.google.com/books?id=8SLUCgAAQBAJ{\&}pgis=1

36. Lawlor DA, Tilling K, Davey Smith G. Triangulation in aetiological epidemiology. International Journal of Epidemiology [Internet]. 2017 Jan;19(R1):dyw314. Available from: https://academic.oup.com/ije/article-lookup/doi/10.1093/ije/dyw314

37. Sterne JAC, Smith GD. Sifting the evidence—what’s wrong with significance tests? BMJ. 2001 322(7280):226–31.

38. Henningsen A, Hamann JD. systemfit: A Package for Estimating Systems of Simultaneous Equations in R. Journal of Statistical Software [Internet]. 2007 Dec;23(4):1–40. Available from: https://www.jstatsoft.org/index.php/jss/article/view/v023i04/v23i04.pdf

39. Steiger JH. Tests for comparing elements of a correlation matrix. Psychological Bulletin. 1980 87(2):245–51.

40. Revelle W. psych: Procedures for Psychological, Psychometric, and Personality Research [Internet], Evanston, Illinois: Northwestern University; 2015. Available from: http://cran.r-project.org/package=psych

41. Shakhbazov K, Powell JE, Hemani G, Henders AK, Martin NG, Visscher PM, et al. Shared genetic control of expression and methylation in peripheral blood. BMC genomics [Internet]. 2016 Jan;17(1):278. Available from: http://bmcgenomics.biomedcentral.com/articles/10.1186/s12864-016-2498-4

42. R Core Team. R: A Language and Environment for Statistical Computing [Internet]. Vienna, Austria: R Foundation for Statistical Computing; 2015. Available from: https://www.r-project.org/

43. Bird A. DNA methylation patterns and epigenetic memory. Genes & development [Internet]. 2002 Jan;16(1):6–21. Available from: http://genesdev.cshlp.org/content/16/1Z6.long

44. Cole DA, Preacher KJ. Manifest Variable Path Analysis: Potentially Serious and Misleading Consequences Due to Uncorrected Measurement Error. Psychological Methods. 2014 19(2):300–15.

45. Bose M, Wu C, Pankow JS, Demerath EW, Bressler J, Fornage M, et al. Evaluation of microarray-based DNA methylation measurement using technical replicates: the Atherosclerosis Risk In Communities (ARIC) Study. BMC Bioinformatics [Internet]. 2014 15(1):312. Available from: http://www.biomedcentral.com/1471-2105/15/312

46. Bryant PA, Smyth GK, Robins-Browne R, Curtis N, Novak J, Sladek R, et al. Technical Variability Is Greater than Biological Variability in a Microarray Experiment but Both Are Outweighed by Changes Induced by Stimulation. Khodursky AB, editor. PLoS ONE [Internet]. 2011 May;6(5):e19556. Available from: http://dx.plos.org/10.1371/journal.pone.0019556

47. Zhu Z, Zhang F, Hu H, Bakshi A, Robinson MR, Powell JE, et al. Integration of summary data from GWAS and eQTL studies predicts complex trait gene targets. Nature Genetics [Internet]. 2016 Mar;48(5):481–7. Available from: http://www.nature.com/doifinder/10.1038/ng.3538

48. Richmond RC, Davey Smith G, Ness AR, Hoed M den, McMahon G, Timpson NJ. Assessing Causality in the Association between Child Adiposity and Physical Activity Levels: A Mendelian Randomization Analysis. Ludwig DS, editor. PLoS Medicine [Internet]. 2014 Mar;11(3):e1001618. Available from: http://dx.plos.org/10.1371/journal.pmed.1001618

49. Mancuso N, Shi H, Goddard P, Kichaev G, Gusev A, Pasaniuc B. Integrating Gene Expression with Summary Association Statistics to Identify Genes Associated with 30 Complex Traits. The American Journal of Human Genetics [Internet]. 2017 Mar;100(3):473–87. Available from: http://linkinghub.elsevier.com/retrieve/pii/S0002929717300320

50. Lee SH, Wray NR. Novel genetic analysis for case-control genome-wide association studies: quantification of power and genomic prediction accuracy. PLoS One. 2013 8(8):e71494.

51. Do R, Willer CJ, Schmidt EM, Sengupta S, Gao C, Peloso GM, et al. Common variants associated with plasma triglycerides and risk for coronary artery disease. Nature Genetics [Internet]. 2013 Oct;45(11):1345–52. Available from: http://www.nature.com/doifinder/10.1038/ng.2795

52. Burgess S, Freitag DF, Khan H, Gorman DN, Thompson SG. Using multivariable Mendelian randomization to disentangle the causal effects of lipid fractions. PloS one [Internet]. 2014 Jan;9(10):e108891. Available from: http://journals.plos.org/plosone/article?id=10.1371/journal.pone.0108891

53. Relton CL, Davey Smith G. Two-step epigenetic Mendelian randomization: a strategy for establishing the causal role of epigenetic processes in pathways to disease. International journal of epidemiology [Internet]. 2012 Feb;41(1):161–76. Available from: http://www.pubmedcentral.nih.gov/articlerender.fcgi?artid=3304531{\&}tool=pmcentrez{\&}rendertype=abstract

54. Varbo A, Benn M, Smith GD, Timpson NJ, Tybjaerg-Hansen A, Nordestgaard BG. Remnant cholesterol, low-density lipoprotein cholesterol, and blood pressure as mediators from obesity to ischemic heart disease. Circulation research [Internet]. 2015 Feb;116(4):665–73. Available from: http://www.ncbi.nlm.nih.gov/pubmed/25411050

55. Burgess S, Daniel RM, Butterworth AS, Thompson SG. Network Mendelian randomization: using genetic variants as instrumental variables to investigate mediation in causal pathways. International journal of epidemiology [Internet]. 2015 Apr;44(2):484–95. Available from: http://www.pubmedcentral.nih.gov/articlerender.fcgi?artid=4469795{\&}tool=pmcentrez{\&}rendertype=abstract

56. Richmond RC, Hemani G, Tilling K, Davey Smith G, Relton CL. Challenges and novel approaches for investigating molecular mediation. Human molecular genetics [Internet]. 2016 Oct;25(R2):R149–56. Available from: http://www.ncbi.nlm.nih.gov/pubmed/27439390 http://www.pubmedcentral.nih.gov/articlerender.fcgi?artid=PMC5036871

